# Microbial Hauberks: Composition and Function of Surface Layer Proteins in Gammaproteobacterial Methanotrophs

**DOI:** 10.1101/2024.07.09.602694

**Authors:** Richard Hamilton, William Gebbie, Chynna Bowman, Alex Mantanona, Marina G. Kalyuzhnaya

**Affiliations:** Biology Department, San Diego State University, San Diego, CA 92182 USA

**Keywords:** methanotrophy, surface (S-)layer, glycoprotein, *Methylotuvimicrobium*, *Methylomicrobium*

## Abstract

Many species of proteobacterial methane-consuming bacteria (methanotrophs) form a hauberk-like envelope represented by a surface (S-) layer protein matrix. While several proteins were predicted to be associated with the cell surface, the composition and function of the hauberk matrix remained elusive. Here we report the identification of the genes encoding the hauberk-forming protein in two gamma-proteobacterial (Type I) methanotrophs, *Methylotuvimicrobium buryatense* 5GB1 (EQU24_15540) and *Methylotuvimicrobium alcaliphilum* 20Z^R^ (MEALZ_0971 and MEALZ_0972). The proteins share 40% AA identity with each other and are distantly related to the RsaA proteins from *Caulobacter crescentus* (20% AA identity).

Deletion of these genes resulted in loss of the characteristic hauberk pattern on the cell surface. A set of transcriptional fusions between the MEALZ_0971 and a superfolder green fluorescent protein (sfGFP) further confirmed its surface localization. The functional roles of the hauberk and cell-surface associated proteins, including MEALZ_0971, MEALZ_0972, EQU24_15540, and a copper-induced CorA protein, were further investigated via gene expression studies and phenotypic tests. The hauberk core protein shows constitutive expression across 18 growth conditions. The *M. alcaliphilum* 20Z^R^Δ0971 showed reduced growth at high salinity, high methanol and metal-limited conditions, suggesting a role in cell-envelope stability and metal scavenging.

Overall, understanding the genetics, composition and cellular functions of the S-layers contributes to our knowledge of methanotroph adaptation to environmental perturbations and opens a promising prospect for (nano)biotechnology applications.

## INTRODUCTION

Surface (S-) layers that consist of proteins or glycoproteins forming lattices covering the cells (i.e., SLPs) have been identified in many prokaryotic organisms spanning both bacteria and archaea [1]. These proteins are abundant proteins synthesized by organisms at a high metabolic cost and, thus, scientists have wanted to characterize their functions and determine their role in specific environments [2]. The predicted functions of SLPs in different bacterial species varies based on whether the organisms are gram positive or negative. Predicted functions include cell adhesion, protection, virulence factors, anchoring sites, receptors for phages, porin sites, and molecular/ ion traps [1–3]. SLPs are unique self-assembling cellular structures with numerous applications in (nano)biotechnology as well as materials and environmental sciences [4]. One of the most characterized SLP in gram-negative bacteria is from *Caulobacter crescentus*. Its S-layer protein was identified to be a 98-kDa protein (*rsaA*) that is transported to the outer membrane through a type 1 secretion system (T1SS) [5]. The *rsaA* protein forms a hexamer (p6 symmetry) anchoring to the outer membrane by liposaccharides and makes up about 30% of total cell protein [6–8].

S-layer matrices have been identified in numerous methane-consuming bacteria (methanotrophs), including alpha and gamma proteobacteria and verrucomicrobia [9–12]. Their S-layers have been shown to form matrixes with p6, p4 or p2 symmetry [9–13]. In *Methylococcus capsulatus*, for example, S-layers are arranged in tetragonal (p4) symmetry [14]. In the haloalkaliphilic *Methylotuvimicrobium* strains, which are gammaproteobacterial methanotrophs, the hauberk-like S-layer envelop formed cup-shaped structures packed in hexagonal (p6) symmetry [11, 15].

It has been suggested that the methanotroph hauberk-like envelope is formed by glycosylated proteins; however, the genetics have remained elusive [11]. Studies to determine dominant outer cell envelope proteins for a haloalkaliphilic *Methylotuvimicrobium* strain (*M. alcaliphilum* 20Z) have revealed a number of proteins including CorA and CorB homologs [16]. The CorA protein has been implicated in copper acquisition in some methanotrophs [16–18]. In *M. album* BG8, a *corA* mutant showed impaired growth on methane [17, 19]. A similar phenotype was observed for *M. alcaliphilum* 20Z^R^ . The *M. alcaliphilum* 20Z^R^ Δ*corA* (MEALZ_2831) mutant was unable to grow on methane but grew on methanol, while the *corB* (MEALZ_2832) mutant grew faster on methane than the wild-type [16]. In addition, it has been shown that the *corA* mutant showed major ultrastructural changes and appeared to contain the S-layer protein within the intercellular space and only loosely attached to the outer cell surface. Thus, it has been suggested that CorA is involved not only in copper acquisition, but also in attaching the S-layer to the outer cell envelope [16]. An outer membrane protein has recently been identified in an acidophilic verrucomicrobial methanotrophic strain *Methylacidiphilum fumariolicum* SolV. The outer membrane protein (WP_009059494) was shown to be highly expressed under a variety of conditions forming a porin with ten antiparallel β-barrels [20]. Considering that S-layers were mostly investigated in extremophilic methanotrophs, their function was often associated with the envelope stability.

In this study, we focus on the identification of genes that encode S-layer proteins in *M. alcaliphilum* 20Z^R^ and *M. buryatense* 5GB1. Both *Methylotuvimicrobium* strains have been shown to possess hauberk-like S-layers with hexagonal cup-like structures. We investigated the composition of their cell-envelope proteins, particularly the matrix core protein, and explored S-layer protein roles in cell envelop stability, metal uptake and methanotrophic metabolism by integrating -omics studies, mutagenesis, and cell-surface imaging. This study solves the decade-long puzzle of the hauberk-protein composition and the role of the surface matrix in gammaproteobacterial methanotrophs while also paving the way to employ the hauberk matrix as functionalized nanomaterials [4].

## RESULTS

### Identification of putative S-layer proteins (SLPs): -omics studies

Initial SLP candidates were identified using existing RNAseq transcriptomics datasets. Reasoning that the SLP must be generated in high amounts and noting that most SLPs are larger than average proteins, the top 50 expressed genes in existing datasets of methane *M. buryatense* 5GB1 and *M. alcaliphilum* 20Z^R^ were sorted based on size [21]. One candidate (MEALZ_0971) in *M. alcaliphilum* 20Z^R^ and its homolog (EQU24_15540) in *M. buryatense* 5GB1 were identified as both highly expressed and greater than 5kb in size in both methanotrophs. The proteins share 40% amino acid (AA) identity. A smaller gene (MEALZ_0972) adjacent to MEALZ_0971 was identified in *M. alcaliphilum* 20Z^R^. Based on transcript mapping data, MEALZ_0972 forms an operon with MEALZ_0971. The small protein does not have a homologue in the genome of *M. buryatense* 5GB1.

We compared the expression levels of MEALZ_0971 and MEALZ_0972 with those of the most abundant cellular proteins, such as particulate methane monooxygenase (pMMO) and the methanol dehydrogenases (MxaF and XoxF) (Figure 1). Transcriptomic datasets for *M. alcaliphilum* 20Z^R^ were analyzed across 18 different environmental conditions that are known to control gene expression levels including +/- copper, calcium vs lanthanides, methanol vs methane, and high to low salinities. Two bioreactor cultures were also evaluated (methane and methanol) of *M. buryatense* 5GB1 [22–24]. A set of proteins were included that were previously identified as associated with the hauberk envelope, including CorA, CorB, MEALZ_0810, and MEALZ_0811 (Figure 1). Only the large proteins (MEALZ_0971 and EQU24_15540) have significantly above average gene expression in both strains. In *M. alcaliphilum* 20Z^R^, the MEALZ_0971 gene product also showed the highest levels of spectral counts in the proteome. The expression of CorA and MEALZ_0810- MEALZ_0811 correlated with copper supplementation, with all genes being induced by copper limitation. The expression of MEALZ_0972 gene was constitutive across most conditions, but significantly lower than the MEALZ_0971 expression.

**Figure 1.**
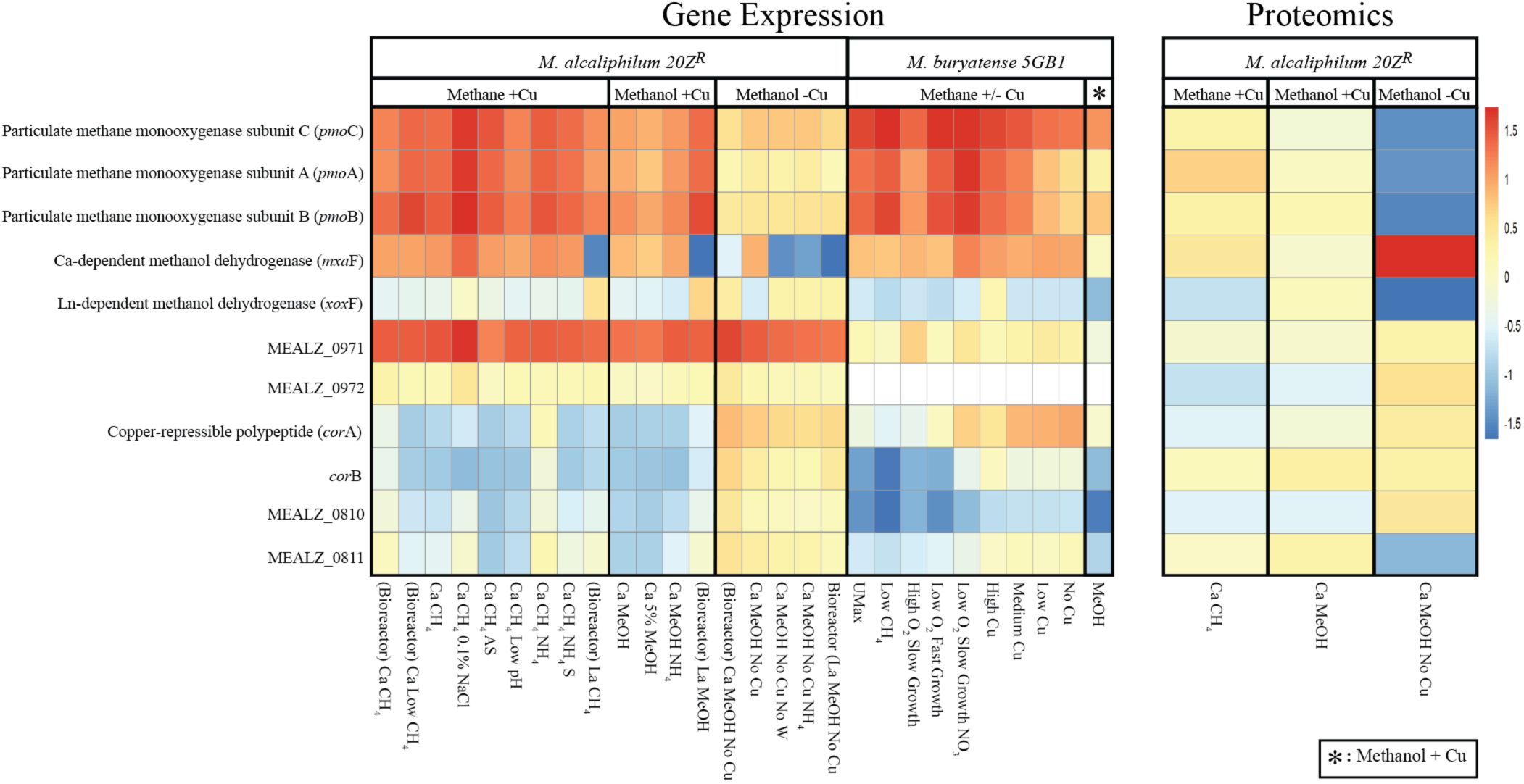
Heatmap of transcriptomic and proteomic expression levels in putative outer membrane protein in *M. alcaliphilum 20Z^R^* and *M. buryatense* 5GB1genes compared to key metabolic enzymes across different growth conditions.

### Comparative genomics

Assessment of the translated amino acid sequence of the predicted methanotrophic SLPs using the multiple alignment program MAFFT showed that they have 20.6% AA identity to the S-layer protein (RsaA) from *Caulobacter crescentus* (Figure 2*). rsaA* lies in the same operon as two of the three type 1 secretion system (TISS) genes, permease/ATPase (*rsaD*) and the periplasmic adapter subunit (*rsaE*). Upstream of both proposed S-layer genes in 20Z^R^ and 5GB1 lies all three genes for the TISS, a common arrangement for S-layer genes, with the permease/ATPase of *M. alcaliphilum* 20Z^R^ having 45% identity and the periplasmic adapter subunit having 34% identity with *Caulobacter Crescentus* (Figure 2) [25, 26]. The calcium-binding domain of the RsaA protein from *Caulobacter* lays in homologous regions of the proposed S-layer proteins for the strains 20Z^R^ and 5GB1 [6, 27]. Additional homologs for the S-layer of *M. alcaliphilum* 20Z^R^ were identified using BLAST in the genomes of other methanotrophs known to have S-layers, *Methylomicrobium album* BG8 (30% AA identity using BLAST and 19% AA identity using MAFFT) and *Methylosarcina lacus* (33% AA identity), as well as in other halophiles or haloalkaliphiles, *Halomonas salina* (48% AA identity), *Desulfonatronum thioautotrophicum* (38% identity), *Desulfonaspira thiodismutans* (33% AA identity), *Oceanicola* sp. HL-35 (33% AA identity), and *Marinobacterium litorale* (29% AA identity). In addition, a homolog was present in *Acidovorax* sp KKS102 (30% AA identity). The correlation of these genes with strains known to have S-layers and their location adjacent to predicted TISS, suggested they were candidates for encoding S-layer proteins.

**Figure 2.**
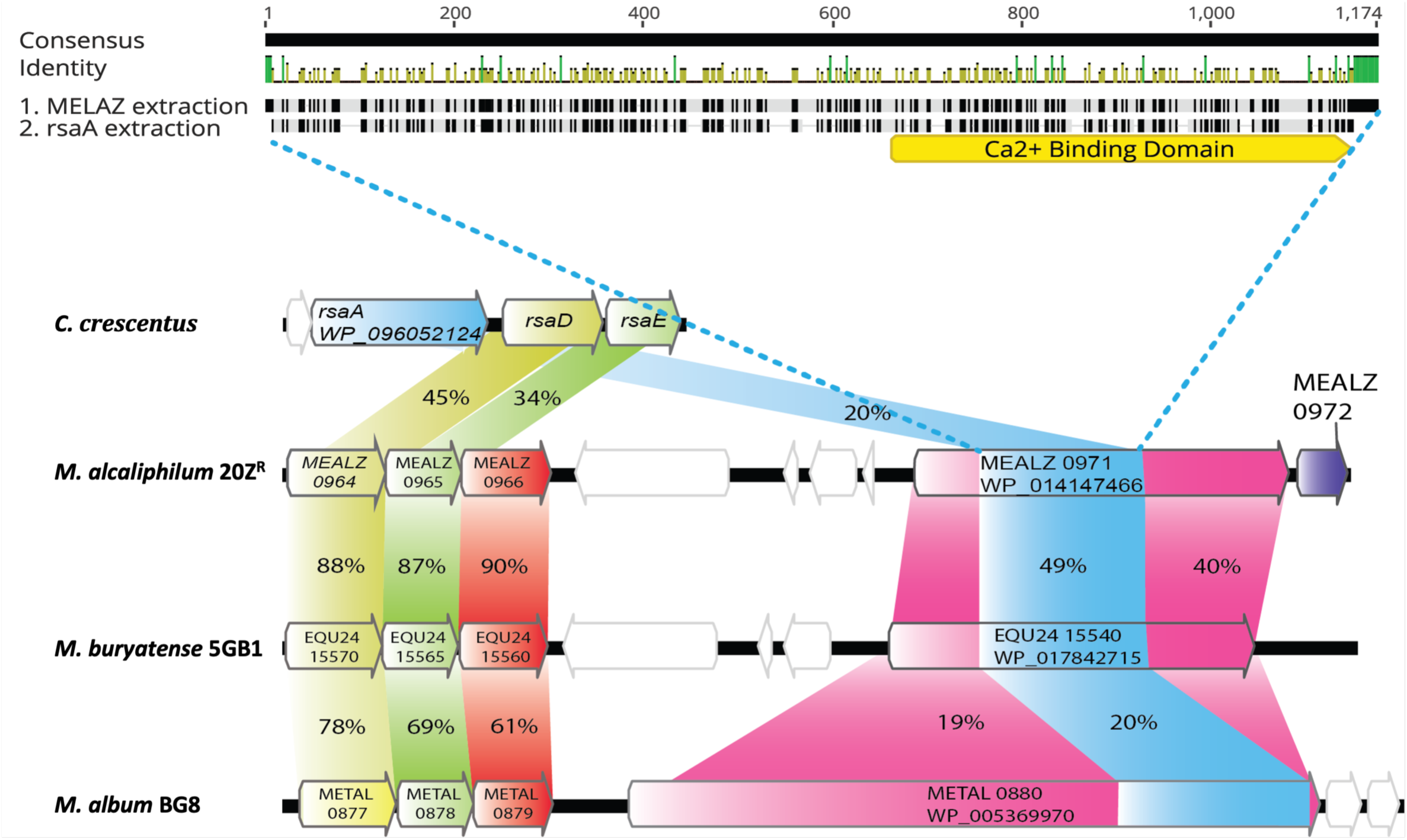
Comparison of identified gene loci in the genome and protein homology in *M. alcaliphilum 20Z^R^* and *M. buryatense 5GB1* to known SLP protein (rsaA) in *C. crescentus* and identified homologue in *M. album BG8.* The homology of RsaA to each predicted SLP protein is highlighted in blue. RsaD and RsaE proteins for T1SS were compared for similarity highlighted in yellow and green. *C. crescentus* is lacking the putative type 1 secretion outer membrane protein which was compared for similarity in 20Z^R^, 5GB1, and BG8 highlighted in red. Lastly MEALZ_0972 highlighted in purple was solely identified in *M. alcaliphilum* 20Z^R^ and was in the same operon as the predicted surface layer protein.

### Protein structure: Alphafold

S-layer proteins are typically attached to lipopolysaccharides (LPS) by non-covalent interactions in most studied gram-negative bacteria [6, 26]. To characterize the MEALZ_0971 protein cell attachment, we compared it to the already characterized cryo-EM S-layer protein (RsaA) from *Caulobacter crescentus.* The N-terminus of the RsaA protein is anchored to LPS on the cell surface of *Caulobacter crescentus* [6]. The RsaA protein forms a hexamer structure with itself with N-terminal regions localized next to each other (Figure 3A). The MEALZ_0971 protein’s homology to the RsaA protein is located toward the C-terminal end (highlighted in gold in Figure 3C). When the translated MEALZ_0971 protein was processed through DeepTMHMM, it was predicted to be anchored with a transmembrane region located near the N-terminal position of the protein (highlighted in purple, Figure 3C), signifying a potential functional difference in their attachment mechanisms.

**Figure 3.**
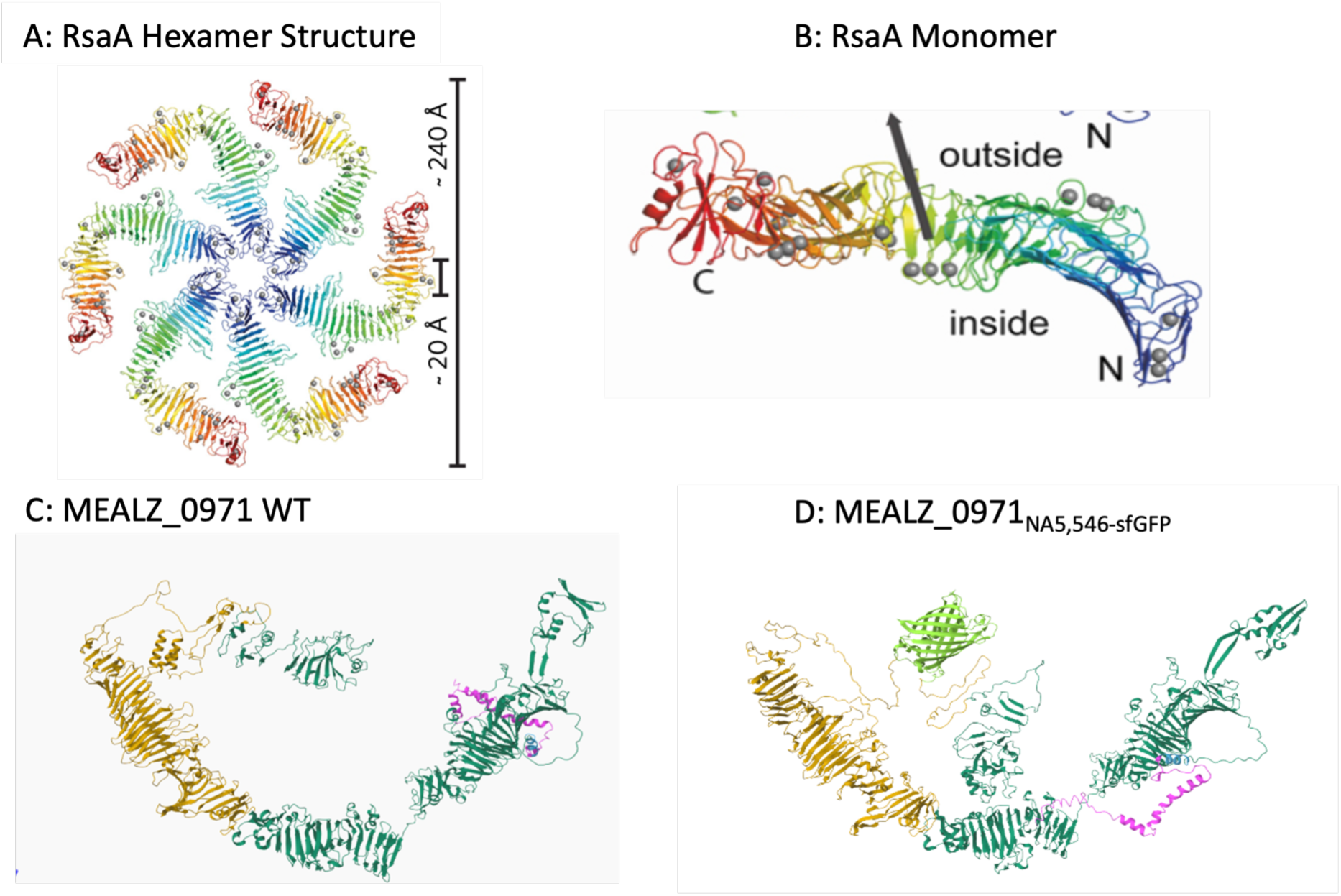
RsaA protein complex from Bharat, Kureisaite-Ciziene, et al. 2017. A: RsaA oriented in Hexamer structure. B: RsaA monomer. Blue -> Red indicates N terminal to C terminal amino acid ordering. The dark blue region represents the domain of the protein that binds to the lipopolysaccharide on the surface of the cell. C: AlphaFold of WT protein. D: AlphaFold of MEALZ_0971 with GFP integrated 1000bps up from C-terminal end of gene. The Gold Regions represent the regions homologous to RsaA (exists on outside of the cell; part of green region). The Green Regions represent the predicted regions existing outside of the cell. The Blue Regions represent the predicted transmembrane regions of the protein. The Pink Regions represent the predicted inner membrane regions of the protein suspected to be in the periplasm. The Light Green Regions in MEALZ_0971 GFP represent the predicted structures of GFP inserted into MEALZ_0971.

### Mutagenesis and Scanning Electron Microscopy (SEM) studies

Single MEALZ_0971 (20Z^R^Δ0971) and MEALZ_0972 (20Z^R^Δ0972) knockout mutants of *M. alcaliphilum* 20Z^R^ were constructed. Single and double knockouts of *M. buryatense* 5GB1 genes EQU24_15540 (5GB1Δ15540) and EQU24_07680 (5GB1Δ*corA*), as well as complemented mutants of 5GB1Δ15540 (5GB1Δ15540::P89 EQU24_15540) were generated and provided for this study by Dr. M. E. Lidstrom (University of Washington). The 5GB1Δ*corA* mutant was included to investigate the protein’s role in S-layer assembly. Once the mutant strains were constructed, they were analyzed by electron microscopy to verify any phenotypic changes to the hexagonal matrix on the surface of the cells. As expected, the scanning electron micrographs of wild type cells of *M. alcaliphilum* 20Z^R^ (Figure 4A) and *M. buryatense* 5GB1 (Figure S1A) verified the S-layer phenotype is present as arrays of hexagonal pits (i.e., the hauberk envelop). In contrast, the *M. alcaliphilum* 20Z^R^Δ0971 and *M. buryatense* 5GB1Δ15540 mutants showed phenotypes with smooth ridges and valleys across the cell surface, which is a characteristic of the outer-membrane (Figure 4A and Figure S1B). The 20Z^R^Δ0972 mutant had some hexagonal pits along the surface but also showed globular structures (i.e., bumps) all over the surface unlike that of 20Z^R^Δ0971 mutant which was mostly smooth (Figure 4A). The 5GB1*corA* mutant cells have cell-surface morphology similar to wild type indicating that the protein is not responsible for the formation of hauberk-like structures (Figure S1C).

**Figure 4.**
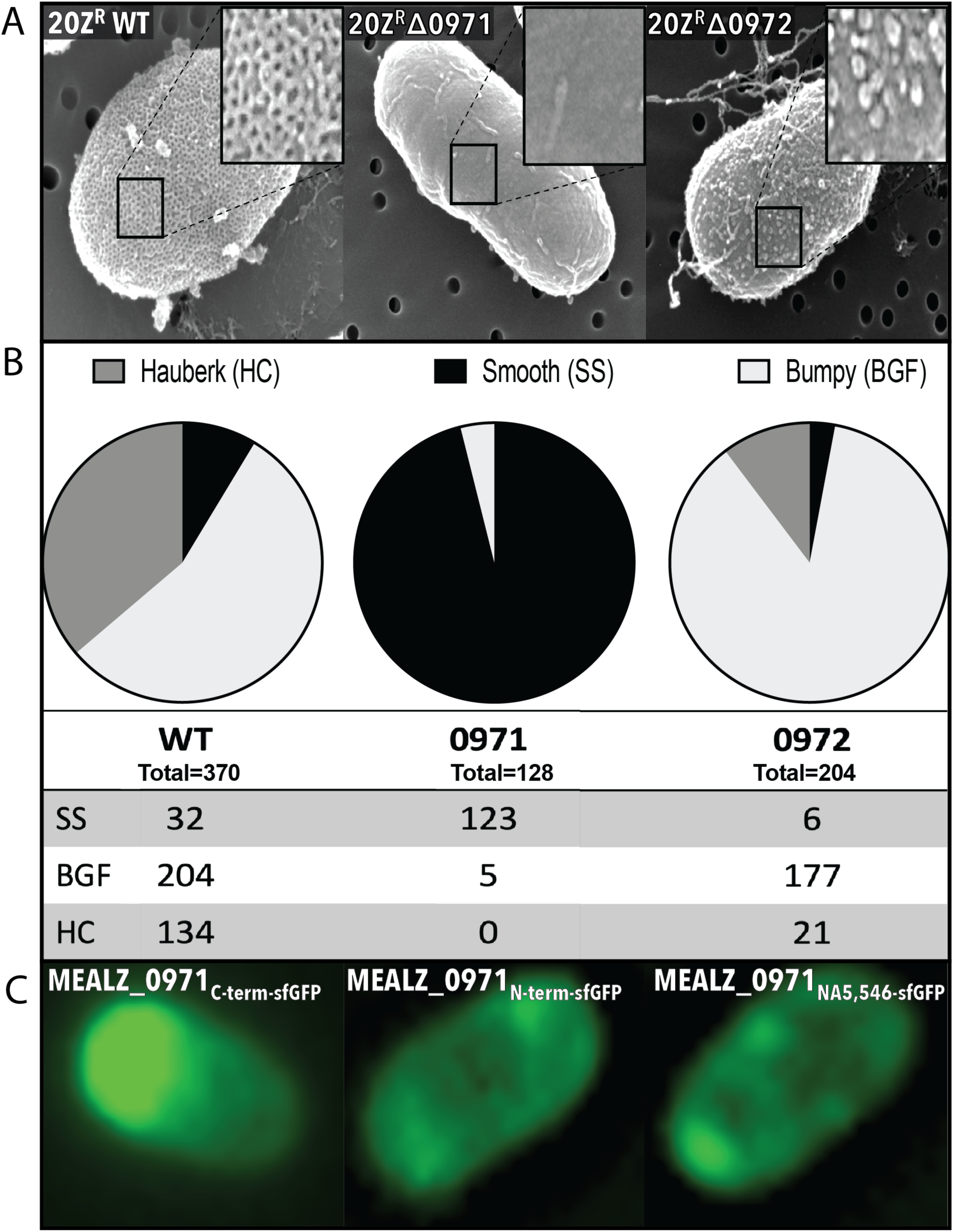
Analysis of phenotypic changes of the surface of the cells with proposed surface layer proteins mutants using both scanning electron microscopy (SEM) and fluorescent microscopy. A: SEM of the SLP on 20Z^R^ WT cells, 20Z^R^ with MEALZ_0971 knocked out, and 20Z^R^ with MEALZ_0972 knocked out. B: Quantification using SEM micrographs of the surface of *M. alcaliphilum 20Z^R^*, *M. alcaliphilum 20Z^R^*Δ0971, and *M. alcaliphilum 20Z^R^*Δ0972. Surface represented as displaying hauberk crystalized SLP, bumpy unorganized SLP, or smooth cell surface. C: Fluorescent microscopy of sfGFP integrated into the SLP at the C-terminal, N-terminal, and at the NA5,546 position from the start codon.

We found that capturing micrographs of individual cells using electron microscopy can be challenging. The hauberk layer can be masked by the presence of glycosylation, the fixation protocol, and/or cell growth stage. Since we observed some smooth-surface cells even among wild type cells, a qualitative assessment of cells morphology was conducted and cells were divided into three main categories: hauberk-covered (HC), bumpy globular features (BGF), or smooth surface (SS). The *M. alcaliphilum* 20Z^R^ wild type presented a total of 134 HC cells, 204 BGF cells, and 32 SS cells (Figure 4B). The number of cells covered by hauberk-like envelop was significantly reduced in *M. alcaliphilum* 20Z^R^Δ0972, which presented a total of 21 HC cells, 177 BGF cells, and 6 SS cells (Figure 4B). We were not able to find any HC cells in the *M. alcaliphilum* 20Z^R^Δ0971 mutant. The cell population of the mutant was represented mostly by SS cells (123) and a few BGF cells (Figure 4B).

### Superfolder green fluorescent protein (sfGFP) tagging and fluorescent microscopy

To further confirm the MEALZ_0971 protein localization, we constructed a set of translational fusions with sfGFP. The fluorescent marker was integrated into MEALZ_0971 of *M.alcaliphilum* 20Z^R^ at the N_term_, C_Term_, and at the NA5546 position from a putative start codon of the protein. The Alphafold-predicted protein demonstrated no change in configuration with sfGFP integrated at the sites (Figure 3D show internal integration). All three strains were investigated using fluorescent microscopy. When the protein was tagged at the N-term, there was little accumulation in the cell; the protein was distributed evenly across the cell surface (Figure 4C). When MEALZ_0971 was sfGFP-tagged internally at the NA5546 site, there was also little accumulation of fluorescent signal within the cell (Figure 4C). Thus, the live-imaging data for the MEALZ_0971_Nterm-sfGFP_ and internal MEALZ_0971_NA5546-sfGFP_ integration suggest that the MEALZ_0971 protein is associated with the *M. alcaliphilum* cell envelop. However, when MEALZ_0971 was tagged at the C-terminal end of the protein, there was an accumulation of sfGFP at the poles of the cell (Figure 4C). This may indicate impaired folding or impaired secretion caused by the GFP integration at the C terminus.

### Phenotyping studies

Given the homology observed in S-layer proteins among halophilic microbes, subsequent testing was conducted to determine their potential role in cell-envelope stability in hypersaline environments. The strains were grown in 1 of 4 conditions (0.5%, 1.5%, 3%, and 6% NaCl) slowly increasing the salinity of the medium. At 3% salinity, the optimal growth condition, *M. alcaliphilum* 20Z^R^ Δ0971 had a higher growth rate (0.052±0.006 h^-1^) compared to wild type and Δ0972 mutant (Figure 5A). The strain also displayed a higher growth rate at low salinity (0.5 and 1.5%); however, when the strains were grown at the highest salinity of 6% NaCl, only the *M. alcaliphilum* 20Z^R^ Δ0971 strain showed a statistically significant decrease in growth rate, which dropped from 0.030±0.001 to 0.021±0.001 h^-1^.

**Figure 5.**
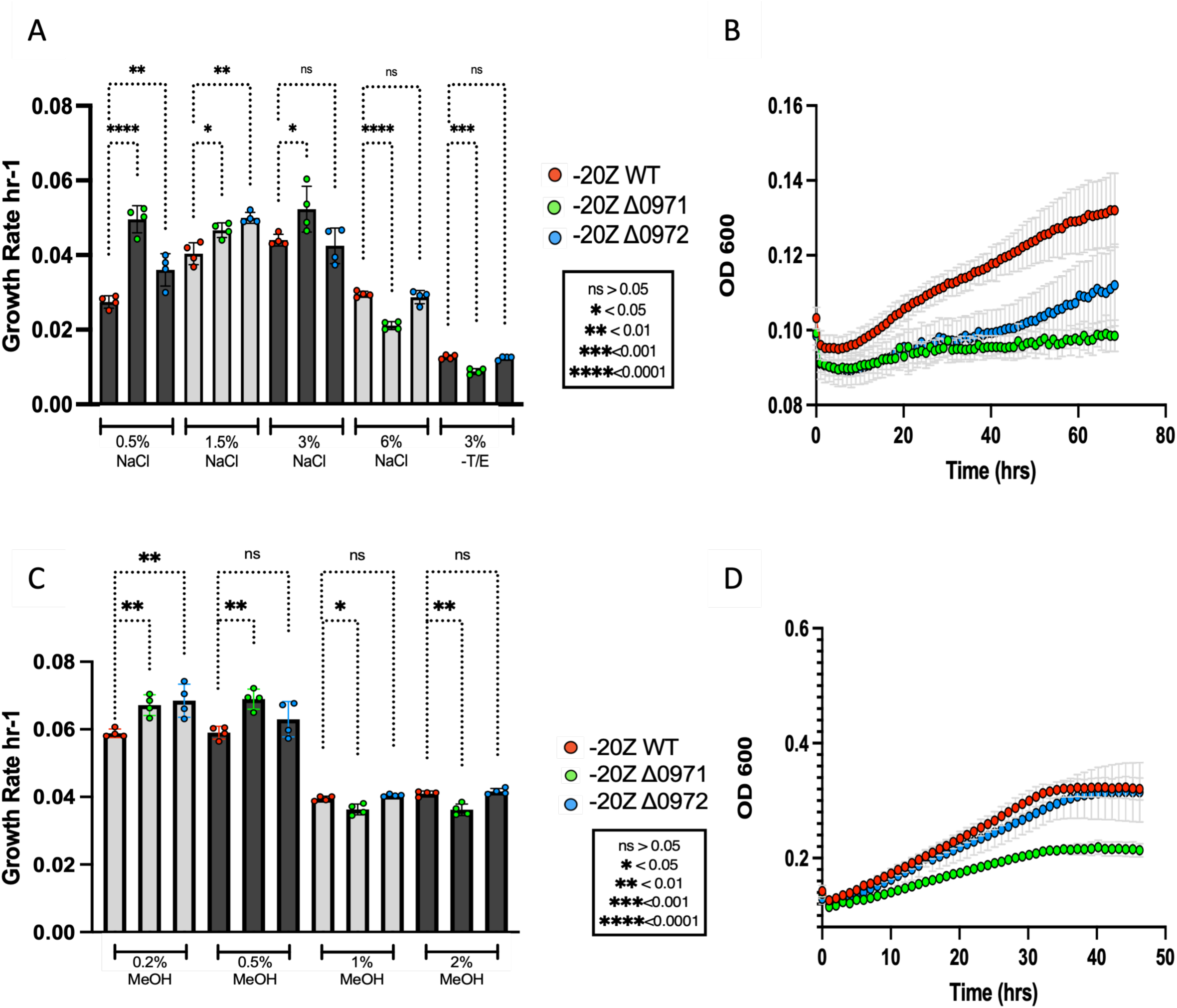
Growth rate of *M.alcaliphilum* 20ZR and ΔMEALZ_0971 and ΔMEALZ_0972 mutants. A. Different salinity (0.5-6% NaCl w/v) or metal limited conditions. B: Growth curve of 20Z^R^ WT, 20Z^R^ Δ0971, and 20Z^R^ Δ0972 grown without T/E. C: Different methanol supplementation (0.2-2% v/v). D: Growth curve of 20Z^R^ WT, 20Z^R^ Δ0971, and 20Z^R^ Δ0972 grown in 6% NaCl.

To test if the MEALZ_0971 and MEALZ_0972 proteins contributor to stress tolerance and membrane stability, the wild type and mutant strains were grown in continually increasing methanol conditions from 0.2% to 2%. The mutant strains of *M. alcaliphilum* 20Z^R^ Δ0971 and Δ0972 showed higher growth rates compared to WT at lower methanol concentrations of 0.2% and 0.5% (Figure 5C). When the methanol concentration was increased to 1%, 20Z^R^ Δ0971 had a significant decrease in growth rate from 0.040±0.001 to 0.036±0.001 h^-1^. When grown in 2% MeOH there was once again a significant decrease in growth rate from 0.041±0.001 to 0.036±0.001. In contrast, the *M. alcaliphilum* 20Z^R^Δ0972 mutant growth rate did not change significantly and was similar to wild type.

To evaluate contribution of the proteins in metal uptake, the wild type and mutant strains of *M. alcaliphilum* were grown under metal-limited conditions. All tests were carried out with the optimal salinity (3%) with 0.2% methanol. However, the growth media was supplemented only with nitrogen (KNO_3_), sulfur (MgSO_4_) and phosphorous sources. When grown in medium lacking minerals, the 20Z^R^ Δ0971 showed a significant decrease in growth rate from the WT strain from 0.012±0.001 to 0.008±0.001 h^-1^ (Figure 5A), while the growth of 20Z^R^ Δ0972 mutant was similar to wild type.

Similar experiments were carried out with constructed mutants for 5GB1 (5GB1Δ*corA* and 5GB1Δ15540. Under the mineral limitation, the mutant 5GB1Δ*corA—*showed a significant decrease in growth rates compared to wild type while the 5GB1Δ15540 showed no difference (Figure S2).

### Inductively coupled plasma (ICP) optical emission spectrometry data

To further test the role of S-layer proteins in metal uptake, the metal compositions of the cell biomass grown at optimal or metal-limited conditions were investigated (Table 1). At optimal conditions, the mutant cells showed higher uptake of Fe and Mn after normalization to account for varieties in biomass, while the rest were at the levels similar to wild type.

**Table 1.**
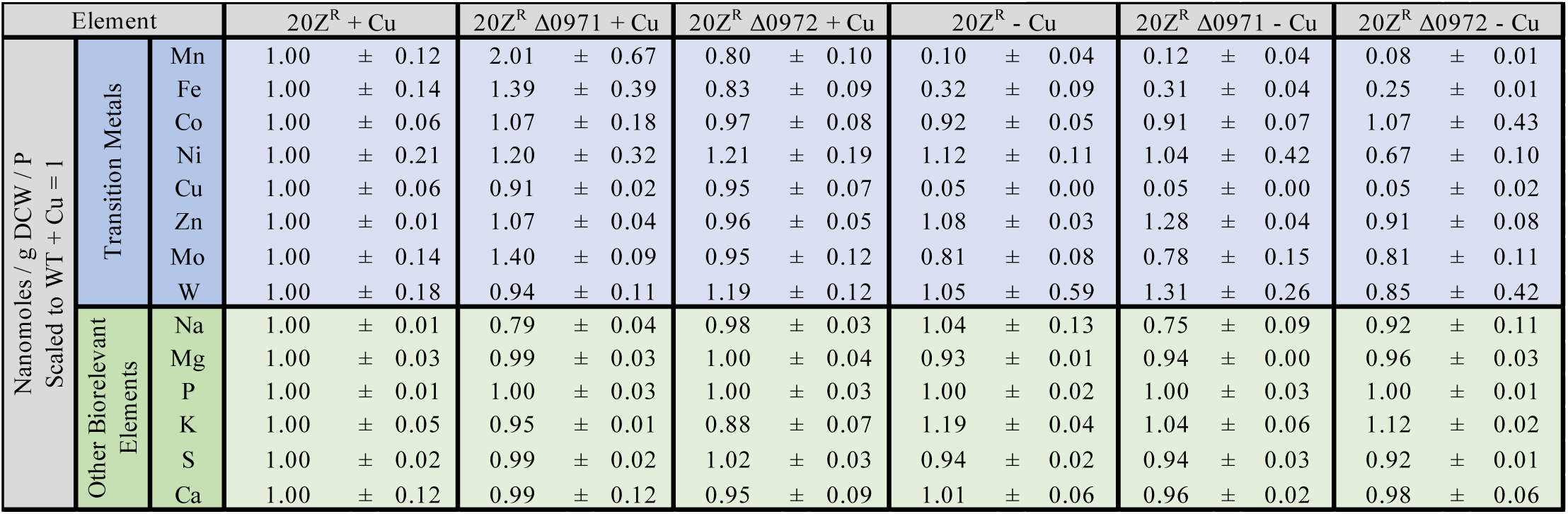
ICP analysis of *M.alcaliphilum* 20Z^R^ WT and mutant cells.

| Element |  |  | 20Z<sup>R</sup> + Cu | 20Z<sup>R</sup> Δ0971 + Cu | 20Z<sup>R</sup> Δ0972 + Cu | 20Z<sup>R</sup> - Cu | 20Z<sup>R</sup> Δ0971 - Cu | 20Z<sup>R</sup> Δ0972 - Cu |
| --- | --- | --- | --- | --- | --- | --- | --- | --- |
| Nanomoles / g DCW / P<br>Scaled to WT + Cu = 1 | Transition Metals | Mn | 1.00 ± 0.12 | 2.01 ± 0.67 | 0.80 ± 0.10 | 0.10 ± 0.04 | 0.12 ± 0.04 | 0.08 ± 0.01 |
|  |  | Fe | 1.00 ± 0.14 | 1.39 ± 0.39 | 0.83 ± 0.09 | 0.32 ± 0.09 | 0.31 ± 0.04 | 0.25 ± 0.01 |
|  |  | Co | 1.00 ± 0.06 | 1.07 ± 0.18 | 0.97 ± 0.08 | 0.92 ± 0.05 | 0.91 ± 0.07 | 1.07 ± 0.43 |
|  |  | Ni | 1.00 ± 0.21 | 1.20 ± 0.32 | 1.21 ± 0.19 | 1.12 ± 0.11 | 1.04 ± 0.42 | 0.67 ± 0.10 |
|  |  | Cu | 1.00 ± 0.06 | 0.91 ± 0.02 | 0.95 ± 0.07 | 0.05 ± 0.00 | 0.05 ± 0.00 | 0.05 ± 0.02 |
|  |  | Zn | 1.00 ± 0.01 | 1.07 ± 0.04 | 0.96 ± 0.05 | 1.08 ± 0.03 | 1.28 ± 0.04 | 0.91 ± 0.08 |
|  |  | Mo | 1.00 ± 0.14 | 1.40 ± 0.09 | 0.95 ± 0.12 | 0.81 ± 0.08 | 0.78 ± 0.15 | 0.81 ± 0.11 |
|  |  | W | 1.00 ± 0.18 | 0.94 ± 0.11 | 1.19 ± 0.12 | 1.05 ± 0.59 | 1.31 ± 0.26 | 0.85 ± 0.42 |
|  | Other Biorelevant Elements | Na | 1.00 ± 0.01 | 0.79 ± 0.04 | 0.98 ± 0.03 | 1.04 ± 0.13 | 0.75 ± 0.09 | 0.92 ± 0.11 |
|  |  | Mg | 1.00 ± 0.03 | 0.99 ± 0.03 | 1.00 ± 0.04 | 0.93 ± 0.01 | 0.94 ± 0.00 | 0.96 ± 0.03 |
|  |  | P | 1.00 ± 0.01 | 1.00 ± 0.03 | 1.00 ± 0.03 | 1.00 ± 0.02 | 1.00 ± 0.03 | 1.00 ± 0.01 |
|  |  | K | 1.00 ± 0.05 | 0.95 ± 0.01 | 0.88 ± 0.07 | 1.19 ± 0.04 | 1.04 ± 0.06 | 1.12 ± 0.02 |
|  |  | S | 1.00 ± 0.02 | 0.99 ± 0.02 | 1.02 ± 0.03 | 0.94 ± 0.02 | 0.94 ± 0.03 | 0.92 ± 0.01 |
|  |  | Ca | 1.00 ± 0.12 | 0.99 ± 0.12 | 0.95 ± 0.09 | 1.01 ± 0.06 | 0.96 ± 0.02 | 0.98 ± 0.06 |

## Discussion

Many species of methanotrophic bacteria form a layer of self-assembled glycosylated proteins on the cell surface. However, the composition and function of additional cell envelope proteins have remained unsolved. This study was designed to identify the genetics of the S-layer matrix in gammaproteobacterial methanotrophs. Previous attempts to identify the matrix using targeted proteomics after polyacrylamide gel electrophoresis (PAGE) failed to detect the core protein. It has been speculated that once assembled, the S-layer matrix does not dissociate even after prolonged heat treatment in the presence of common detergents, like SDS, and reagents such as DTT or beta-mercaptoethanol [20]. In this study we used gene expression and non-targeted proteomics data to identify the most highly expressed genes in two methanotrophic bacteria. The initial hypotheses were (1) since the proteins form an envelope wrapped around the cell, they must be highly abundant and represented by a large protein; and (2) since the extracellular proteins are the most likely target for predator attachment (i.e., viruses), they are under a strong selection pressure and should show little conservation among homologs from two closely related strains, such as 5GB1 and 20Z^R^. We identified two proteins that fit all criteria—the MEALZ_0971 gene in *M. alcaliphilum* 20Z^R^ and the EQU24_15540 gene in *M. buryatense* 5GB1. The genes are highly expressed and share 40% amino acid identity.

The comparative genomic studies showed that the proteins encoded by MEALZ_0971 and EQU24_15540 genes share some homology (up to 20%) with well characterized surface layer protein (RsaA) in *C. crescentus*. The SEM images demonstrate that the deletion of MEALZ_0971 and EQU24_15540 leads to elimination of hauberks structures from the surface of the cells, indicating that the proteins contribute to the S-matrix formation. The fluorescent microscopy imaging of two cellular populations with translational sfGFP fusions at the N_ter_ and mid-position of the MEALZ_0971 protein, showed its cell-surface localization. The Z-stack images suggest that the protein is forming an envelope that coats the cells. Based on the data we suggest that MEALZ_0971 and EQU24_15540 encode alkaliphilic methanotroph S-layer core proteins.

One of the translational fusions, MEALZ_0971_Cter-sfGFP_ provided initial insights into the secretion mechanism. When MEALZ_0971 was tagged at the C-terminal loci, secretion was disrupted, supporting the idea that it is being secreted by the type 1 secretion system (T1SS) since proteins transported through T1SS have C-terminal recognition sites [28]. T1SS secretion genes are part of the *rsaA-*operon in *C. crescentus*, and the system has been shown to secrete the S-layer protein [5]. Gene homologs of all three components of T1SS secretion were found upstream of MEALZ_0971 and EQU24_15540. It is tempting to speculate that the genes encode the S-layer protein secretion machinery; however, this needs to be validated by additional studies.

Once the genetics of the S-layers were identified, the next task was to gain insight into functional significance in methanotrophs. Our data showed that MEALZ_0971 and possibly MEALZ_0972 proteins stabilize the cell envelope at high salinities and high methanol conditions. Furthermore, the mutants displayed reduced growth in in metal-limited environments, suggesting a role in metal uptake. To further interrogate the role of the MEALZ_0971 and MEALZ_0972 proteins in metal uptake, we investigated metal content of wild type and mutant cell biomass through ICP-OES. No significant changes were observed, with the exception of somewhat higher accumulation of iron and manganese in the mutants.

The deletion of *corA*, that encodes a putative copper-binding and S-layer associated protein, did not impact the assembly of hauberk-like structures on cell surface in *M. buryatense* 5GB1. Furthermore, the *M. buryatense* 5GB1Δ*corA* grew normally on methane (data not shown).

The 5GB1Δ*corA* phenotype contradicted the phenotypes of *M. alcaliphilum* 20Z^R^Δ*corA* and *M. album* BG8 Δ*corA*, which lost the ability to grow under methane and have a weakened association of S-layers with the cell surface [16–17]. The omics data strongly suggest that CorA and CorB are regulated by copper availability. However, the lack of this copper binding enzyme does not impact the growth of *M. buryatense* 5GB1 in the same way as it does to *M. alcaliphilum* 20Z^R^, most likely because contrary to 20Z^R^, the 5GB1 strain also possesses the iron-dependent soluble methane monooxygenase system (sMMO) and, thus, it can grow without copper under methane growth conditions [29].

We also found a few intriguing correlations—the growth phenotype of the 5GB1Δ15540 followed the phenotype of CorA mutant. The CorA protein has been suggested to contribute to copper uptake. We suggest that the SLP scaffold acts as a dock for other proteins involved in metal uptake. Additional studies are underway to confirm this function of the S-layer matrix.

Overall, it appears that methanotrophic SLPs do not significantly impact cell growth under optimal growth conditions; however, they become more essential under different types of membrane stability stressors. Considering that membrane stability and transport functions are interconnected, further investigations are needed to better understand the contribution of SLPs to metal transport.

## MATERIALS AND METHODS

### Strains and growth conditions

*M. buryatense* 5GB1C and *M. alcaliphilum* 20Z^R^ strains were grown at 30°C with methane (25% headspace) or 0.2% methanol on a nitrate mineral salts medium with 0.75% NaCl for 5GB1 and 0.3% NaCl for 20Z^R^ at pH 9 [33]. For all 20Z^R^ phenotyping studies the strains were grown in 48-well plates in a SpectraMax iD5 Multi-Mode Microplate Reader. Cultures were first grown in specified medium in a 48-well plate until they reached exponential growth around an optical density of 0.3-0.6. These cultures were then used to inoculate the microplate wells with fresh media to an initial OD of 0.1 (total volume 400µl). The setting in the plate reader was kept at a temperature of 30°C with continuously shaking on high in orbital motions. Optical density readings were taken every hour for 3 days or until cultures reached stationary phase of growth. For 5GB1 phenotyping studies the strains were grown in 125 mL shake flasks (with 25mL media). Cultures were first grown in specified medium till they reached expolnential growth of 0.3-0.6 and inoculated into fresh medium to an initial OD of 0.1 (total colume 25mL).

The following *Escherichia coli* strains were used for cloning and gene transfer experiments: *E.coli* S17-1 (lab stock), *E.coli* DH5α (New England Biolabs (NEBlab), Catalog Number C2987H, *fhuA2Δ(argF-lacZ)U169 phoA glnV44 Φ80Δ(lacZ)M15 gyrA96 recA1 relA1 endA1 thi-1 hsdR17*) and E.coli DH10B (NEBlabs, Catalog Number C3019H: *Δ(ara-leu) 7697 araD139 fhuA ΔlacX74 galK16 galE15 e14-ϕ80dlacZΔM15 recA1 relA1 endA1 nupG rpsL (Str^R^) rph spoT1 Δ(mrr-hsdRMS-mcrBC)*). *E.coli* strains were grown using LB broth or LB agar media (Miller, Difco BD Life Sciences) and incubated at 37°C. The following antibiotics were applied when required: kanamycin (100µg/ml), rifamycin (50µg/ml).

### Sequence Alignment and Comparison

The genomic sequences used for this study were *M. alcaliphilum* 20Z^R^ (FO082060.1), *M. buryatense* 5GB1 (CP035467.1), and *C. crescentus* (CP023315.3). The type 1 secretion system proteins and proposed S-layer proteins were compared to *C. crescentus* using the MAFFT online tool [34, 35]. The outputs were then processed into Geneious to construct the comparative diagram [36].

### Alphafold and Structural Comparison Methods

Alphafold 2 (accessed with Tamarind Bio https://www.tamarind.bio/) was used to predict the protein structures of MEALZ_0971 and MEALZ_0972 with an insertion of superfolder green fluorescent protein (sfGFP) 1000 bp upstream of the C-terminal end of the protein [37, 38]. UniProt BLAST was used to find the RsaA homologous amino acid region in MEALZ_0971 [39]. DeepTMHMM v1.0.24 was used to predict cellular locations (cytosol/periplasm, transmembrane, or extracellular) of amino acids in MEALZ_0971 [40]. Protein Data Bank (PDB) files were visualized and the predicted homologous RsaA, cytosolic/periplasmic, transmembrane, and extracellular amino acid regions were colored with Mol* from RCSB PDB [41].

### Transcriptomics Expression Comparison Heatmap Methods

59 RNA-seq raw reads from Akberdin et al., 2018, Vasquez, 2023, and unpublished studies from the C1-Lab at SDSU representing 27 different growth conditions varying in salinity, pH, nitrogen sources, carbon sources, and metalloenzyme cofactors such as copper (Cu), calcium (Ca), lanthanides (Lns), and tungsten (W) were used in this comparative gene expression study [23, 42]. Raw RNA-seq reads were preprocessed with fastp [43]. Transcript quantification was then performed with Salmon in mapping-based mode with the validateMappings and automatic library type detection arguments against the transcriptome of *M. alcaliphilum* 20Z^R^ (Genbank Accession: FO082060.1 & FO082061.1) [44]. The variance Stabilizing Transformation method from DESeq2 was used to normalize raw transcript counts [45]. The gene expression heatmap was generated using the R pheatmap v1.0.21 package [46].

### Mutagenesis

The mutant strains of *M. buryatense* – 5GB1Δ*corA* (EQU24_07680) and 5GB1Δ15540 were kindly provided by Dr. M.E. Lidstrom (University of Washington). The mutants of *M.alcaliphilum* 20Z^R^ were generated using conjugation with pCM433 vector as described previously [29, 33, 47]. The upstream and downstream flanking regions of the target gene were amplified by PCR using Thermo Scientific Phusion Flash PCR Master Mix. These flanking regions were ligated into the linear form of the pCM433 backbone using NEBuilder^®^ HiFi DNA Assembly Master Mix and transformed into NEB 10-beta competent *E. coli* cells. Colonies were tested by colony PCR as described [33]. Positive constructs were then transformed into *E. coli* S17-1 cells used for bi-parental conjugation with *M.alcaliphilum* 20Z^R^. The strain then underwent through selection on kanamycin followed by counterselection with sucrose to knockout the vector. Colonies were then tested for knockout of each gene by PCR and validated by sequencing.

### Translational fusions of MEALZ_0971 with sfGFP

To tag MEALZ_0971 with superfolded green fluorescent protein (sfGFP), the pCM433 allelic exchange vector was used [33]. The sfGFP was codon optimized to 20Z^R^ based on its codon usage bias (CUB) table previously identified [48]. Vectors were constructed with flanking regions of the N-term, C-term, or position 5,546bps to insert sfGFP into each location of the MEALZ_0971. Positive mutants were identified, the tagged gene was amplified using PCR and correct integration was confirmed by sequencing.

### Imaging studies: Scanning electron microscopy (SEM)

The imaging studies were carried out as previously described (53), with following modification: a fixative solution (2.5% glutaraldehyde, 0.1 M cacodylate buffer (CB) was supplemented with 0.5% NaCl to maintain osmolarity during initial fixation. Samples were imaged at the SDSU Electron Microscopy Facility on an FEI Quanta FEG 450 at 20 kV accelerating voltage at a working distance of ∼10 mm.

### Imaging studies: Fluorescent microscopy

Samples were prepared for fluorescent microscopy by growing cells to exponential phase and collecting cells through centrifugation. The cells were then fixed in 4% formaldehyde for 1hour and washed 3X with NMS medium. Samples were mounted and images on an all-in-one Kenence fluorescent microscope BZ-X Series.

### Inductively coupled plasma (ICP) optical emission spectrometry

Cell pellets (n = 3 for each strain) were collected via centrifugation. Each sample (about 20 mg dry cell weight, DCW) was digested in concentrated (68-70%) ultrapure nitric acid at 65 °C for 2-24 hours (until all solids were completely dissolved). Samples were then diluted to 5 mL with the addition of 4.5 mL of ultrapure distilled water. Analysis was performed using an Agilent 5800 ICP-OES with an SPS 4 Autosampler. Three replicates of 1.5 mL per sample were analyzed and the averaged value was used to determine concentrations. Read time was set to 5 seconds, RF power was set to 1.4 kW, stabilization time was 15 seconds, the nebulizer flow was set to 0.70 L/min, the plasma flow 12.0 L/min, Aux flow at 1.00 L/min and the make-up flow set to 0. Measurements were done using the axial viewing mode. Wavelengths were chosen to minimize spectral and chemical interference, and the Agilent software performed peak-fitting to minimize spectral interference. Where interference was high, multiple wavelengths were chosen and averaged to get the reported concentration.

## ACKNOWLEDGEMENTS

The authors thank Dr. Lidstrom for providing the *M. buryatense* mutants, and her insightful comments on the manuscript. We also thank Dr. Ingrid Niesman (SDSU EM Facility) for her knowledgeable advice on the SEM imaging protocol.

